## Supplementary Materials for "Predicting Math Ability using Working Memory, Number Sense, and Neurophysiology in Children and Adults"

**Results of Resting-State Theta and Alpha Activity**

We computed the theta (4-7Hz) and alpha (8-13Hz) peak power calculated with the parameterization method. First, this was done for the frontal-central area (electrodes Fz and FCz). The results regarding the theta and alpha activity are visualized in **Figure S1** and **S2.** Theta and alpha activity in the frontal-central areas calculated with the parameterization method are not related to working memory, number sense or mathematical ability for children and adults.

Secondly, we computed theta and alpha activity over the bilateral parietal-temporal area (electrodes T7, P7, T8, and P8). The results regarding the theta and alpha activity are visualized in **Figure S3** and **S4.** Theta and alpha activity in the bilateral parietal-temporal area calculated with the parameterization method was not related to working memory, number sense or mathematical ability for children and adults.

Thirdly, we investigated the left parietal-occipital (electrodes PO9, P3, and O1) area. The results regarding the theta and alpha activity is visualized in **Figure S5** and **S6.** Theta activity in the left parietal-occipital area was significantly related with mathematical ability (path C: $\beta = -.21$, $SE = .09$, $p = .02$), addition and multiplication problems (path C: $\beta = -.21$, $SE = .09$, $p = .02$), and subtraction and division problems (path C: $\beta = -.18$, $SE = .09$, $p = .04$). This was independent of working memory, number sense and mathematical ability. Alpha activity in the left parietal-occipital area calculated with the parameterization method was not related to working memory, number sense or mathematical ability for children and adults.

Lastly, we looked at theta and alpha activity over the bilateral parietal-occipital area (electrodes PO9, P3, O1, PO10, P4, and O2). The results regarding the theta and alpha activity are visualized in **Figure S7** and **S8.** Theta and alpha activity in the bilateral parietal-occipital area calculated with the parameterization method was not related to working memory, number sense or mathematical ability for children and adults.

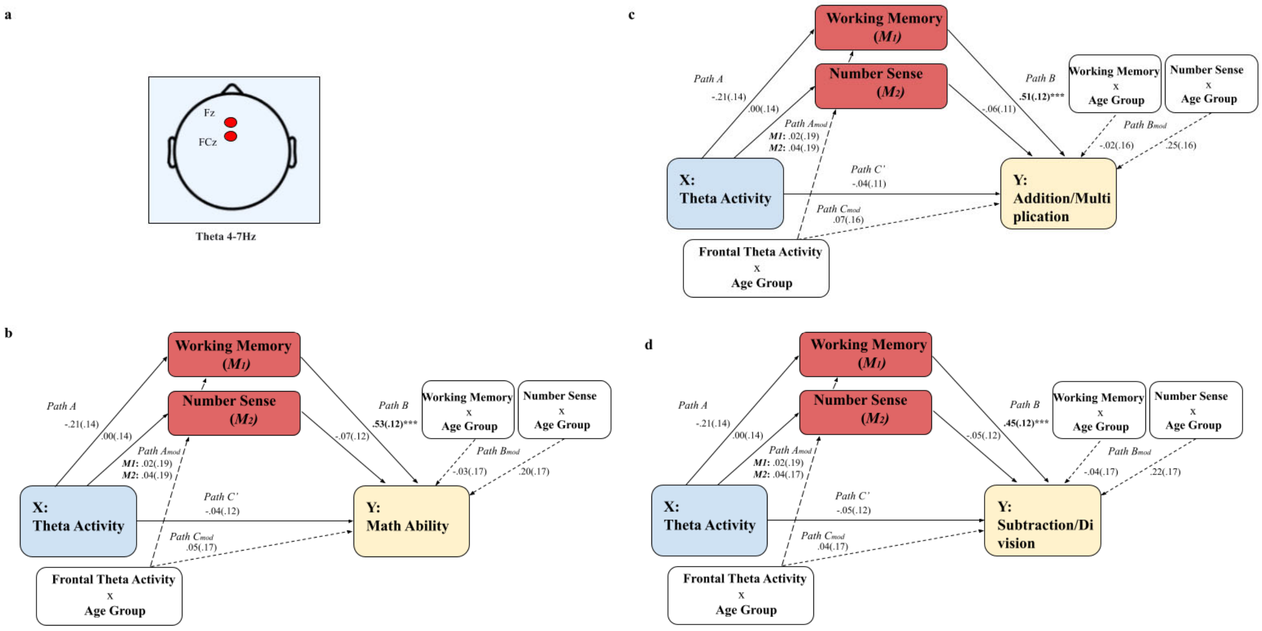


**Figure S1. Moderated mediation model for frontal-central theta activity (n = 105). (a)** Location of the clustered electrodes for the frontal-central area Fz and FCz. **(b)** Moderated mediation model that includes the frontal-central theta activity calculated with the parameterization method and mathematical ability. **(b)** Moderated mediation model that includes the frontal-central theta activity and mathematical addition and multiplication problems. **(c)** Moderated mediation model that includes frontal-central theta activity and subtraction and division problems.***$p < .001$.

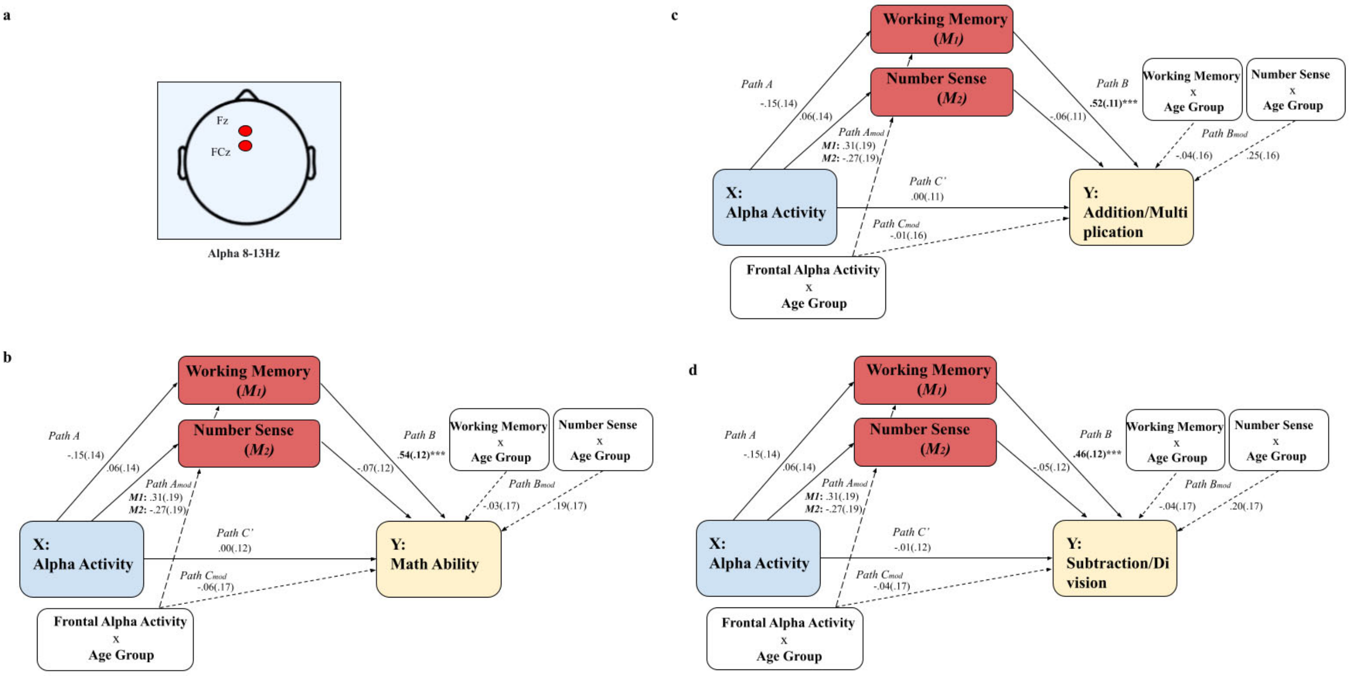


**Figure S2. Moderated mediation model for frontal-central alpha activity (n = 105). (a)** Location of the clustered electrodes for the frontal-central area Fz and FCz. **(b)** Moderated mediation model that includes the frontal-central alpha activity calculated with the parameterization method and mathematical ability. **(c)** Moderated mediation model that includes the frontal-central alpha activity and addition and multiplication problem. **(d)** Moderated mediation model that includes frontal-central alpha activity and subtraction and division problems.***$p < .001$.

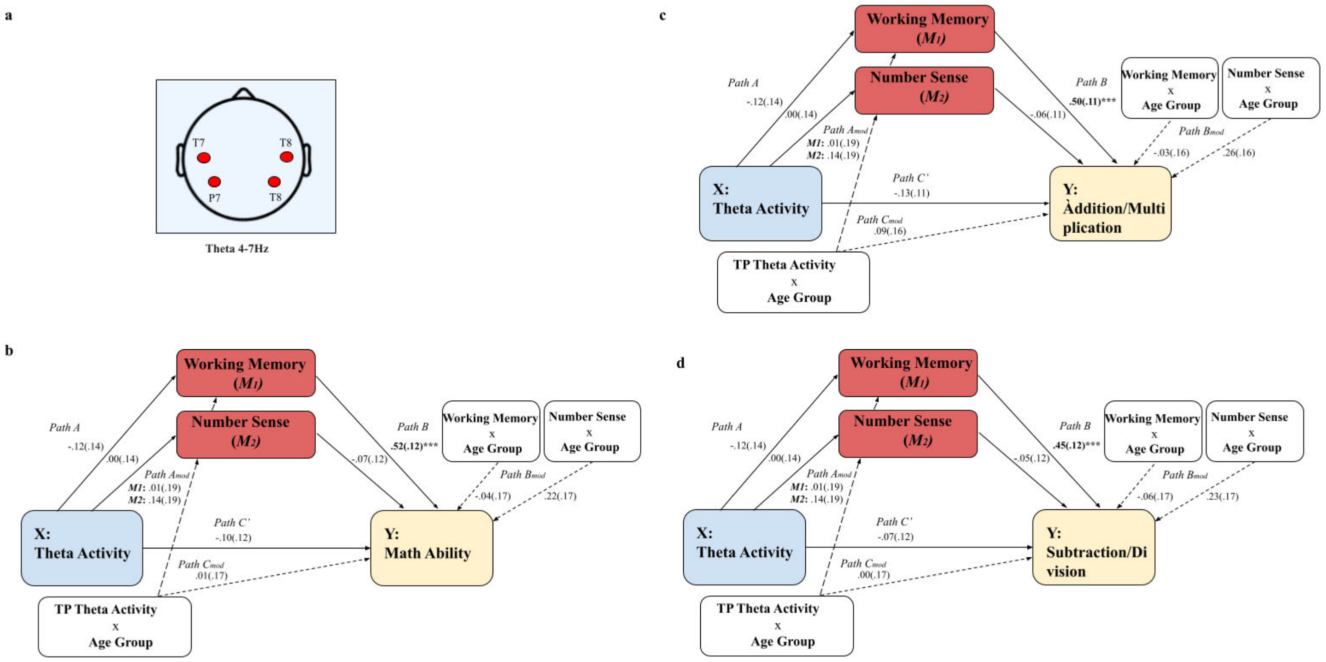


**Figure S3. Moderated mediation model for bilateral parietal-temporal theta activity (n = 105). (a)** Location of the clustered electrodes for the bilateral parietal-temporal area T7, P8, T8, and P8. **(b)** Moderated mediation model that includes the bilateral parietal-temporal theta activity calculated with the parameterization method and mathematical ability. **(c)** Moderated mediation model that includes the bilateral parietal-temporal theta activity and addition and multiplication problems. **(d)** Moderated mediation model that includes bilateral parietal-temporal theta activity and subtraction and division problems.***$p < .001$.

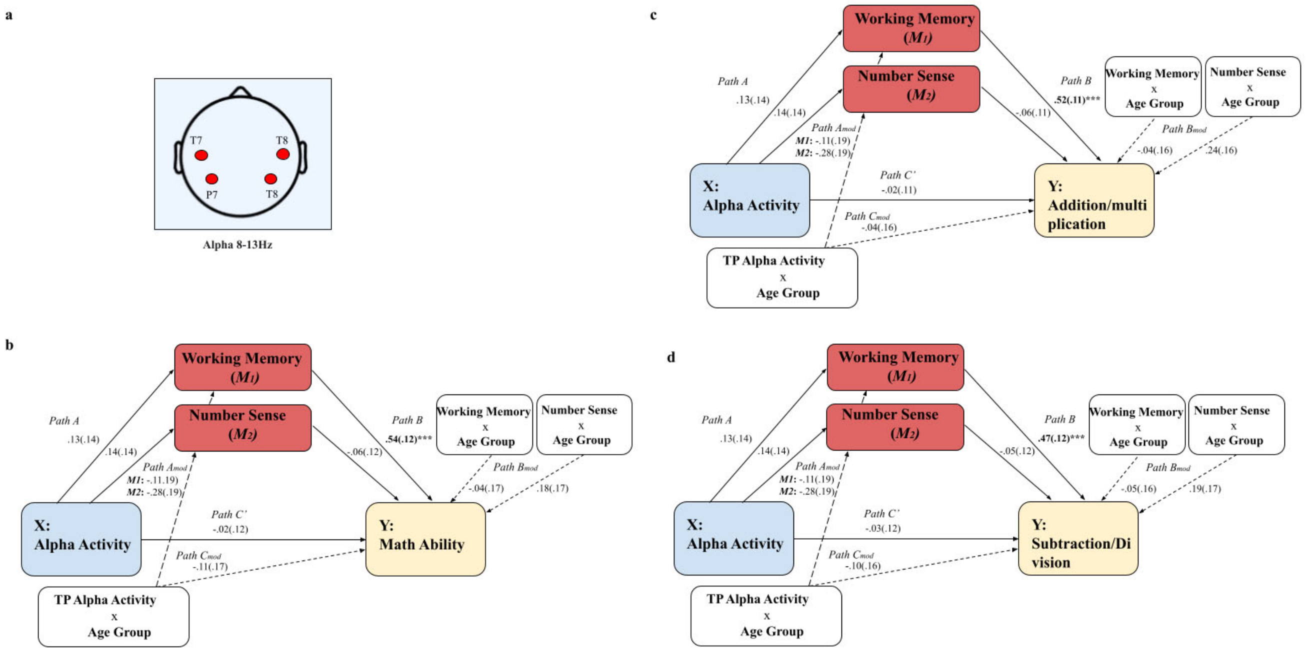


**Figure S4. Moderated mediation model for bilateral parietal-temporal alpha activity (n = 105). (a)** Location of the clustered electrodes for the bilateral parietal-temporal area T7, P8, T8, and P8. **(b)** Moderated mediation model that includes the bilateral parietal-temporal alpha activity calculated with the parameterization method and mathematical ability. **(c)** Moderated mediation model that includes the bilateral parietal-temporal alpha activity and addition and multiplication problems. **(d)** Moderated mediation model that includes bilateral parietal-temporal alpha activity and mathematical subtraction and division problems. ***$p < .001$.

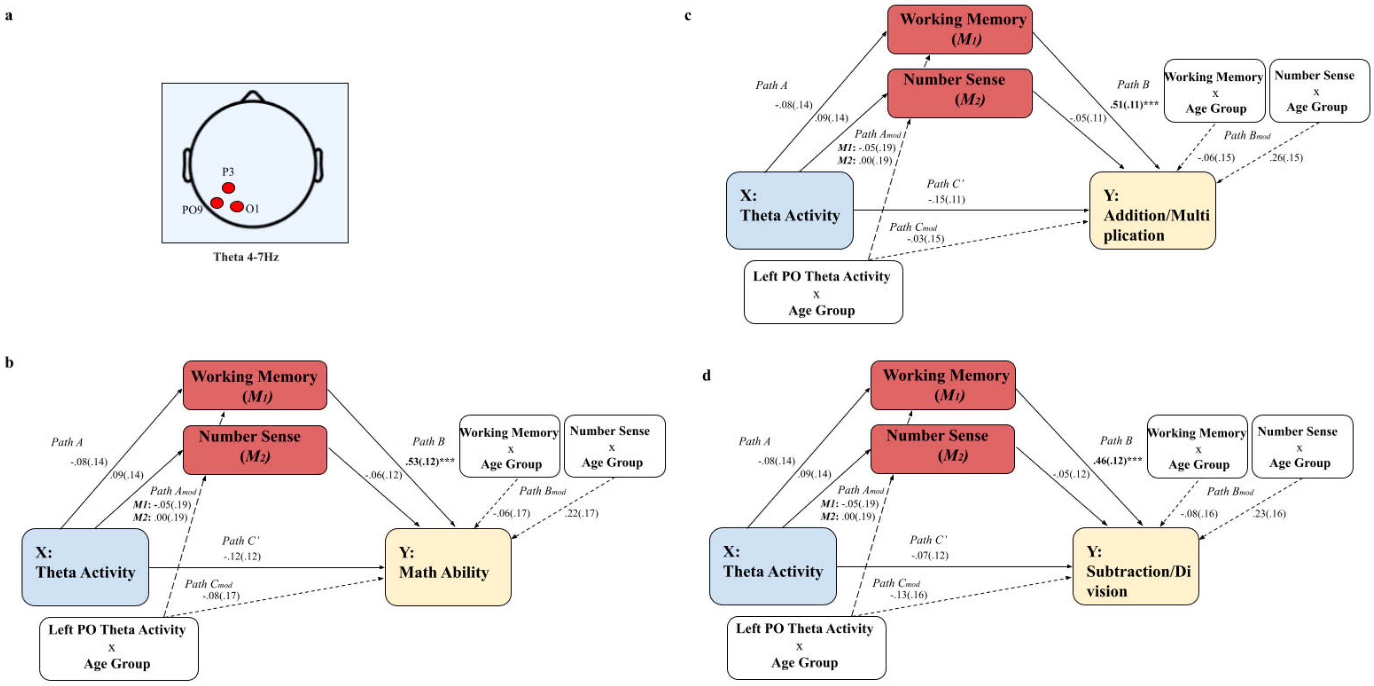


**Figure S5. Moderated mediation model for left parietal-occipital theta activity (n = 105).** **(a)** Location of the clustered electrodes for the left parietal-occipital area P3, O1, and PO9. **(b)** Moderated mediation model that includes the left parietal-occipital theta activity calculated with the parameterization method and mathematical ability. **(c)** Moderated mediation model that includes the left parietal-occipital theta activity and addition and multiplication problems. **(d)** Moderated mediation model that includes left parietal-occipital theta activity and subtraction and division problems. $^*p < .05$, $^{***}p < .001$.

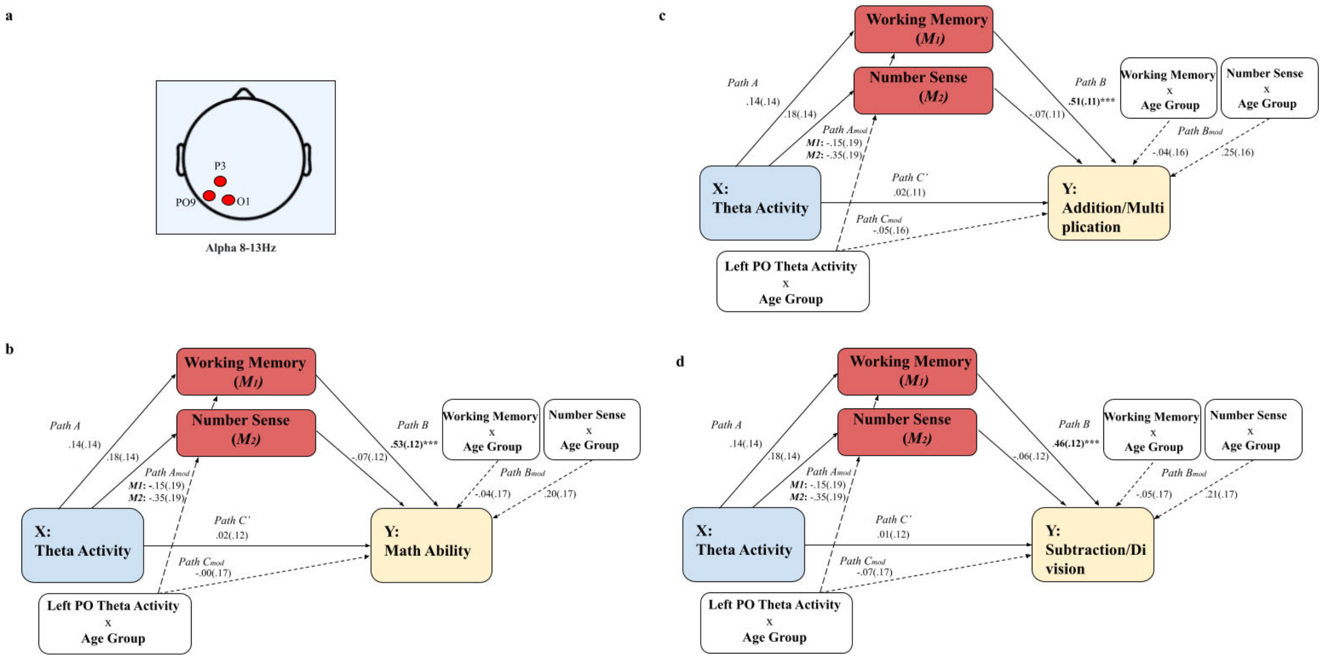


**Figure S6. Moderated mediation model for left parietal-occipital alpha activity (n = 105).** **(a)** Location of the clustered electrodes for the left parietal-occipital area P3, O1, and PO9. **(b)** Moderated mediation model that includes the left parietal-occipital alpha activity calculated with the parameterization method and mathematical ability. **(c)** Moderated mediation model that includes the left parietal-occipital alpha activity addition and multiplication problems. **(d)** Moderated mediation model that includes left parietal-occipital alpha activity and subtraction and division problems.***$p < .001$.

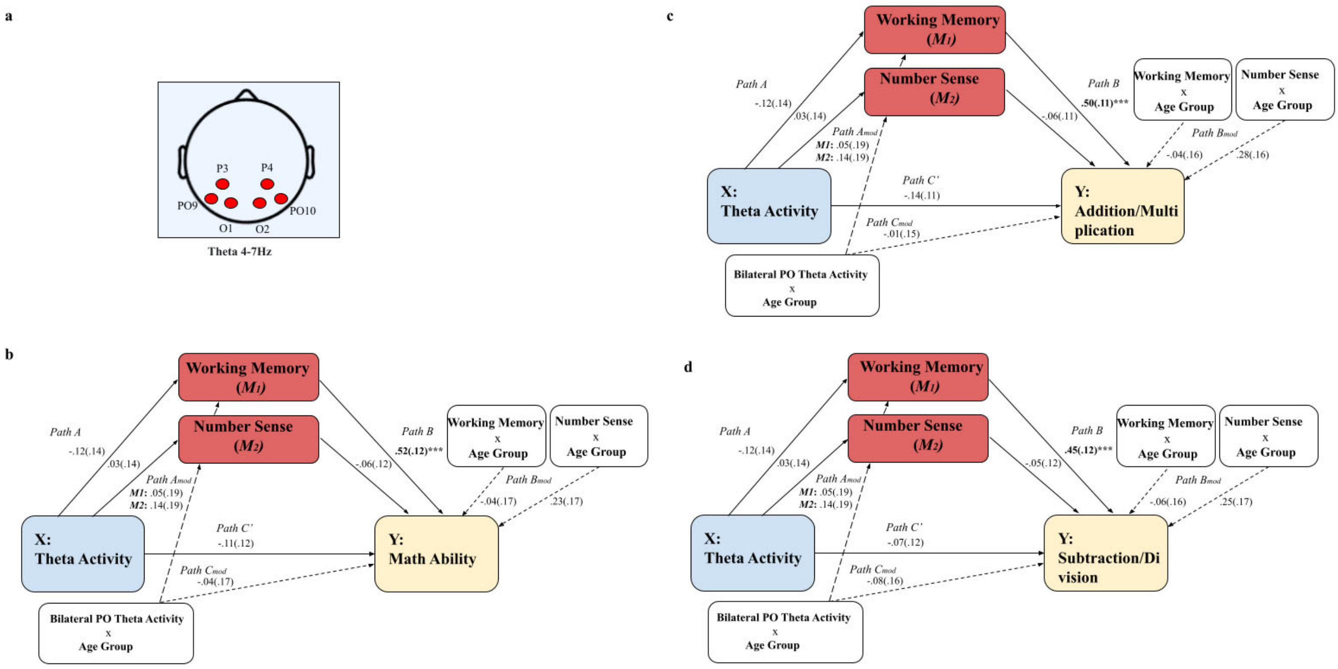


**Figure S7. Moderated mediation model for bilateral parietal-occipital theta activity (n = 105). (a)** Location of the clustered electrodes for the bilateral parietal-occipital area PO9, P3, O1, PO10, P4, and O2. **(b)** Moderated mediation model that includes the bilateral parietal-occipital theta activity calculated with the parameterization method and mathematical ability. **(c)** Moderated mediation model that includes the bilateral parietal-occipital theta activity and addition and multiplication problems. **(d)** Moderated mediation model that includes bilateral parietal-occipital theta activity and subtraction and division problems.***$p < .001$.

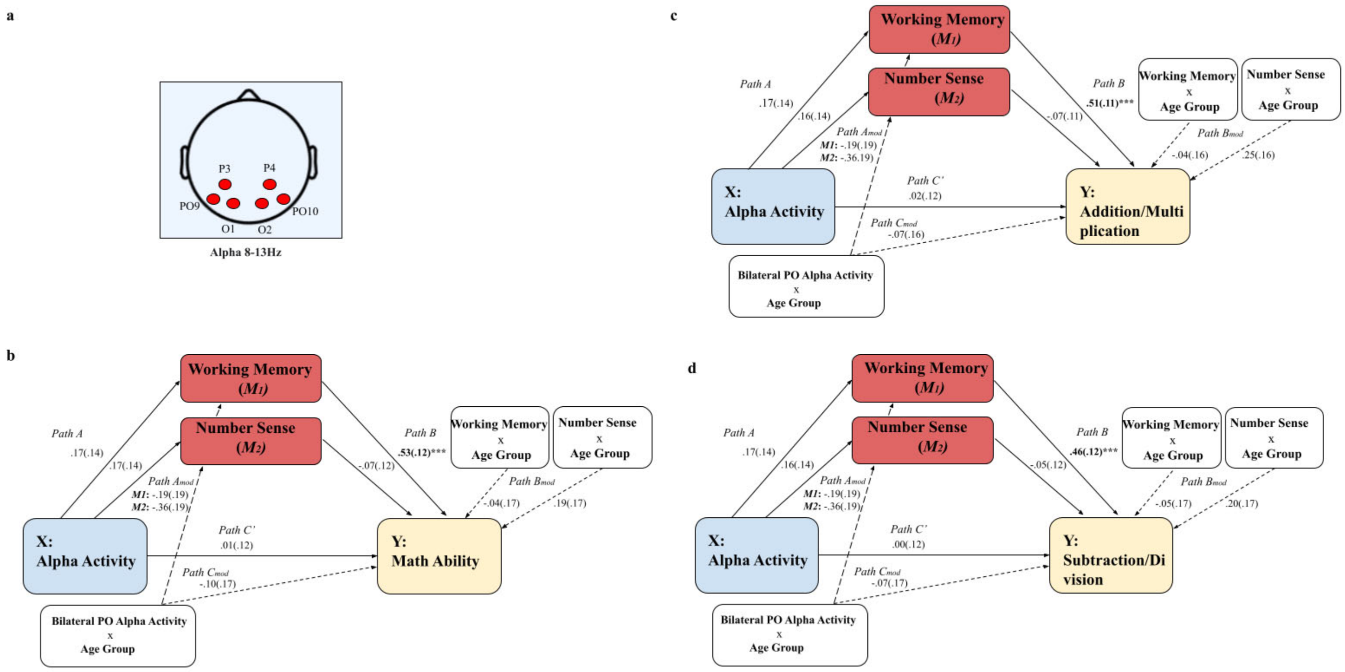


**Figure S8. Moderated mediation model for bilateral parietal-occipital alpha activity (n = 105). (a)** Location of the clustered electrodes for the bilateral parietal-occipital area PO9, P3, O1, PO10, P4, and O2. **(b)** Moderated mediation model that includes the bilateral parietal-occipital alpha activity calculated with the parameterization method and mathematical ability. **(c)** Moderated mediation model that includes the bilateral parietal-occipital alpha activity and mathematical retrieval (addition and multiplication). **(c)** Moderated mediation model that includes bilateral parietal-occipital alpha activity and mathematical procedural calculation (subtraction and division).****p* < .001.
